# Learning Complex Representations from Spatial Phase Statistics of Natural Scenes

**DOI:** 10.1101/112813

**Authors:** HaDi MaBouDi, Krishna Subramani, Hamid Soltanian-Zadeh, Shun-ichi Amari, Hideaki Shimazaki

## Abstract

Natural scenes contain higher-order statistical structures that can be encoded in their spatial phase information. Nevertheless, little progress has been made in modeling phase information of images, and understanding efficient representation of the image phases in the brain. In order to capture spatial phase structure under the efficient coding hypothesis, here we introduce a generative model of natural scenes by assuming independent source signals in a complex domain and non-uniform phase priors for the complex signals. Parameters of the proposed model are then estimated under the maximum-likelihood principle. This approach extends existing methods of independent component analysis for complex-valued signals to the one that utilizes phase information. Using simulated data, we demonstrate that the proposed model outperforms conventional models with a uniform phase prior in blind source separation of complex-valued signals. We then apply the proposed model to natural scenes in the Fourier domain. Real and imaginary parts of the learned complex features exhibit a pair of Gabor-like filters in quadratic phase structure with a similar shape. The proposed model significantly improved the goodness-of-the-fit from the model with a uniform phase prior, indicating that the structured spatial phases are important for removing redundancy in natural scenes. These results predict the presence of phase sensitive complex cells in the visual cortex.

## 1. Introduction

One of the successful guiding principles to understand visual systems in the brain is the efficient coding hypothesis [1]. According to this hypothesis, organizations of a visual system are adapted to regularities in natural scenes that an animal encounters. The efficient coding hypothesis has successfully guided us to construct physiologically plausible statistical models of neurons in early visual cortices [2, 3]. However, most of the previous models extracted information contained only in amplitudes of an image in the Fourier domain, and were blind to its phase structure. Contrary to the assumption of the previous models, it is well known that the phase of an image contains significantly more perceptual information than the amplitude of an image [4]. Perceptually salient features such as edges and bars are encoded in the ventral visual cortex based on their phase congruency [5, 6, 7, 8]. It has also been shown that some complex cells in macaque primary visual cortex,V1, are sensitive to the image phases [9, 10, 11, 12]. Nevertheless, constructing models of a visual system that utilizes the characteristic phase information in the natural scenes remains to be a challenging problem.

The classical linear generative models represent the natural images by linearly combining features (i.e., projection fields), and by weighting them using different coefficients [2, 13]. The coefficients of the features are learned from natural images so that they become as independent as possible, according to the efficient coding hypothesis. Nevertheless it is known that their dependency cannot completely removed. It was pointed out that the residual dependency in the coefficients is conveniently described by using scalar and circular components [14, 15, 16]. This suggests to use complex representation (a pair of real and imaginary features) of the natural images [15, 17, 18]. A successful complex representation model may explain why nearby simple cells in the primary visual cortex are phase quadratic [19, 20], and support psycho-physical studies which suggested image phases may be detected by combining responses of simple cells possessing two odd and even symmetric features [19].

In this study, we present a linear generative model of complex representation of natural images using a superposition of complex features (a pair of features). While we consider independent priors for the amplitude and phase of the coefficients, our attention is particularly paid to the phase distribution. MaBouDi et al. previously demonstrated that local phases of natural scenes detected by Gabor filters are characterized by not only uniform but also non-uniform phase distributions [16]. Motivated by this finding, we model the phase distribution using a mixture of von Mises distributions, and provide inference algorithms under the maximum likelihood principle. We then demonstrate the utility of this approach by both blind source separation of simulated data and analysis of natural scenes.

## 2. Complex-valued Independent Component Analysis

In this section, we present a complex-valued independent component analysis (ICA) that performs blind source separation of complex-valued data, assuming specific statistical structure on their amplitudes and phases.

Let *X* be a column vector of dimension *N* with its elements being complex values 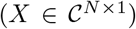. Following the conventional real-valued ICA model, the complex ICA aims to find independent complex sources *S* that satisfies

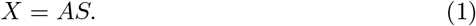

Here 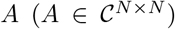 is a complex mixing matrix whose columns represent complex basis functions (features). 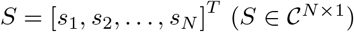 is complex source signals. They are complex coefficients of the complex basis functions in the generative model defined for complex-valued signals. Each element of *S*, namely *s*_*i*_, is given by 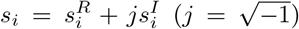, where 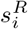 and 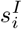 are the real and imaginary components of the coefficient. This equation is also written as *S* = *WX*, where *W* = *A*^−1^ is called a de-mixing matrix. In the following, we provide a model that assumes structured amplitude and phase for the independent source complex signals.

We assume that a sample is generated by the complex linear model using the complex coefficients that are sampled from an independent distribution of *S*, namely 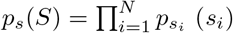. Using the de-mixing matrix *W*, the Jacobian of the above linear transformation is given as det 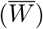 [21]. 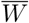 is defined as,

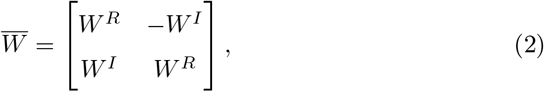

where *W^R^* and *W^I^* are the real and imaginary component of *W*, respectively. Thus given *p*_*s*_(*S*), the probability density function of *X* is obtained as,

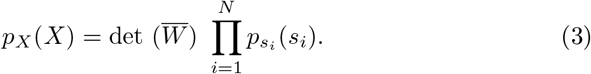

The complex ICA aims to infer the basis matrix *A* (or its inverse *W*) and source signals *S* under the assumption of their independence.

In this study, we propose to model each complex coefficient by polar coordinates, and impose independence between the amplitude and phase components. Namely, we rewrite the complex coefficients as 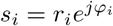, where *r*_*i*_ = |*s*_*i*_| and 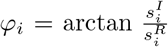 are the amplitude and phase components of *s*_*i*_, respectively. Then the model of the probability density function of *s*_*i*_ is given as

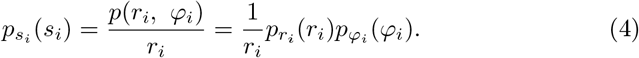

Throughout this paper, we assume that the amplitude distribution 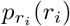 follows the gamma distribution with the shape parameter being 2,

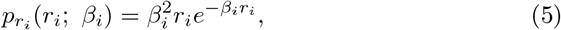

where *β*_*i*_ > 0 is a scale parameter. This distribution resembles the amplitude distribution obtained from responses of complex Gabor filters to natural scenes [16], and imposes sparseness on the complex coefficients. We let the shape parameter be 2 because we found that the optimization algorithm to estimate the complex features under the maximum likelihood principle results in the algorithms proposed by previous studies (see below). While the previous studies utilized only amplitude information (a flat phase prior), our approach based on the maximum likelihood principle allows us to use different types of phase priors, and compare their performance. In the following sections, we derive optimization algorithms for two types of prior distributions for the phases of the complex coefficients of the complex basis functions.

### Circular complex-valued ICA (circular cICA)

In this approach termed a circular cICA (circular cICA), we assume a uniform phase distribution, 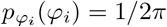. The goal is to estimate the linear transformation *W* such that the elements of the complex coefficient vector, *S*, are as independent as possible through an iterative optimization procedure. We estimate the parameters of the circular cICA model under the maximum likelihood principle. Let ***X***^**obs**^ = (*X*^1^, *X*^2^, …, *X*^*T*^) be *T* complex-valued samples whose dimension is *N*. Given that the samples are independent, Eq. 3 gives the log-likelihood function of the model parameters,

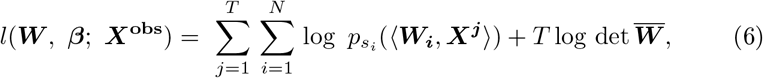

where (***W***_***i***_, ***X***^***j***^) is the dot product between the *i*-th column of ***W*** and ***X***^***j***^. By considering 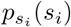 from Eq. 4 with prior knowledge of the amplitude distribution (Eq. 5) and a uniform phase distribution, we obtain

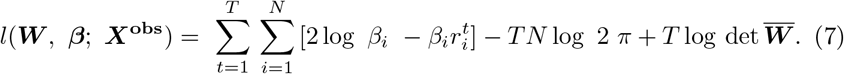

Note that this log-likelihood function generalizes the cost function used in the Fast complex ICA (Fast cICA) [22]. Moreover, since this model assumes a uniform phase distribution for the complex random variables, it is applicable for separation of circular complex random variables [23]. The maximum likelihood estimates (MLEs) of the model parameters can be obtained by gradient ascent algorithms in the complex domain, following the Wirtinger calculus. The Wirtinger calculus was briefly summarized in Appendix 1. The final expressions for the gradients are given as

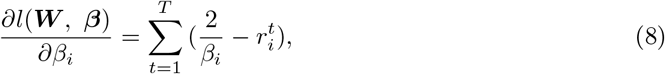

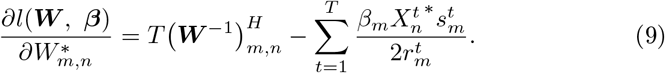

where * displays the conjugate of a complex variable and *H* denotes the hermitian (conjugate transpose) of a complex matrix. See Appendix 2 for the detailed derivation of the gradients.

### Phase-aware complex-valued ICA (phase-aware cICA)

The circular cICA model does not use phase information that may be contained in data such as image patches. For example, it was previously reported that higher-order statistics of natural scene is additionally encoded in non-uniform bimodal phase distributions [16]. In this section, we extend the complex ICA, and construct a model that utilizes the non-uniform phase distributions.

In this approach, we model the phase distribution by a mixture of uniform and von-Mises distributions (see one example of the mixture model of phase distribution in Fig. 1):

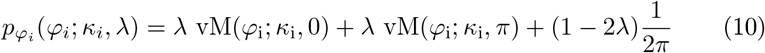

**Figure 1:**
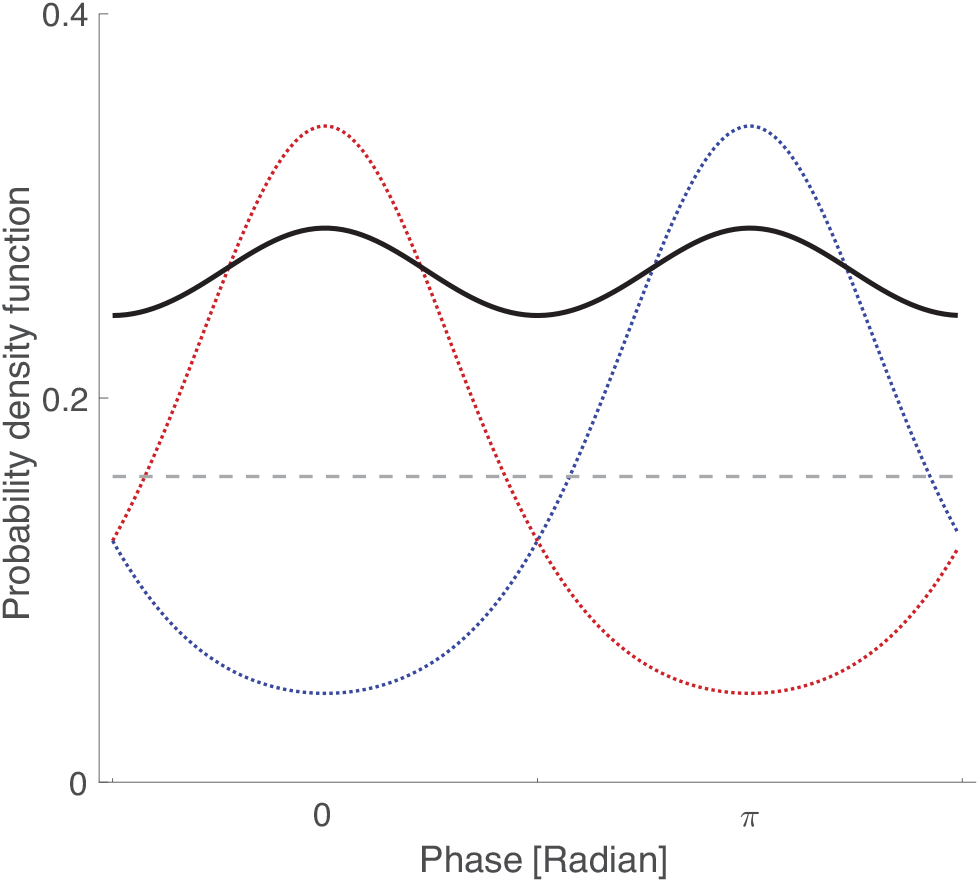
A mixture model of phase distribution. The red and blue dotted curves exhibit two von Mises distributions with the same concentration parameter *κ*, but with different peaks at *µ* = 0 and *µ* = *π*. The grey dashed line shows the uniform distribution. The black curve represents a mixture model of phase distribution that is composed of these three distributions.

Here vM(*φ*_i_; *κ*_i_, 0) is a von-Mises distribution for a circular variable *φ*_*i*_ with zero mean and a concentration parameter *κ*_*i*_. The von-Mises distribution with mean *µ*_*i*_ is defined as,

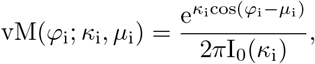

where *I*_0_(.) is the Bessel function of order 0. Given the observations of spatial phase distributions in natural scenes [16], we consider symmetric bimodal phase distributions with two peaks separated by *π*. Here we assume that the peaks of the phase distribution are located at 0 and *π* because the peak phase location is redundant when the features are learned from the data. Note that this model becomes the uniform phase distribution if *λ* = 0 or *κ*_*i*_ = 0 and non-uniform if *λ* ≠ 0 and *κ*_*i*_ ≠ 0. For simplicity we further assume equal contributions from each component (*λ* = 1/3). Then the phase distribution that can cover a uniform and a spectrum of bimodal phase distributions is simplified as

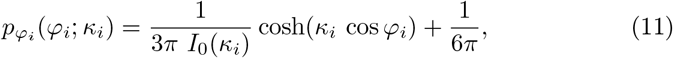

This model of the structured phase distribution incorporates additional information regarding the phase of source signals as opposed to the the previous circular cICA. Thus we call our proposed model the ‘phase-aware complex-valued ICA’ (phase-aware cICA) model. Note that this model becomes the circular cICA with uniform phase distribution when *κ*_*i*_ = 0.

The log-likelihood function of the phase-aware cICA model is

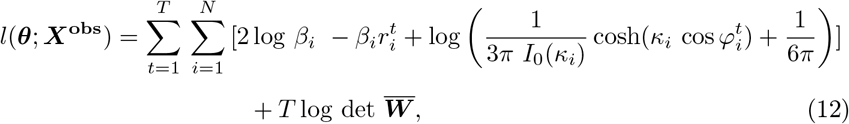

where ***θ*** = (***W***, ***κ***, ***β***) is the set of model parameters. The gradients of the likelihood function of the phase-aware model with respect to the model parameters are derived analytically in Appendix 3, and the final expressions are given as

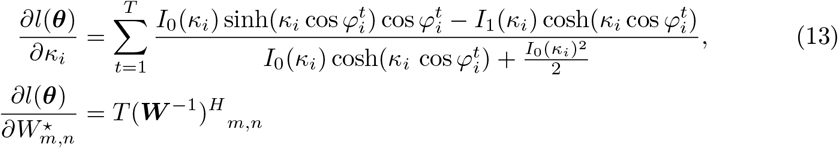

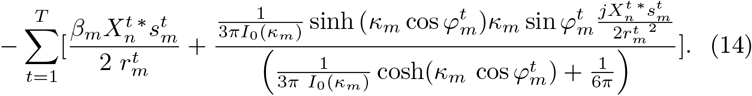

Here, *I*_1_(.) is the Bessel function of order 1. For *β*, the gradient is the same as in Eq. 8.

## 3. Results

### Performance comparison using simulated data

In this section, we evaluate efficiency of source separation by variants of the cICA’s, using an artificially generated dataset. For this goal, we generated a dataset *X* by mixing 10 independent complex-valued source signals with a gamma-distributed amplitude and uniform or non-uniform phase *S*, using a random invertible mixing matrix *A*. We considered two different types of the complex-valued source signals using the polar coordinates: In one data set, phases were sampled from a uniform distribution; in the other data set, phases were sampled from a bimodal distribution (Eq. 10). Amplitudes were sampled from Eq. 5 in both cases. In total, we generated 20 data sets for each case. The mixing matrix and parameters of the probability density function of sources were chosen randomly for each data set.

We then applied three cICA methods to estimate the de-mixing matrix *W*. We evaluated their performance as follows:

1. A set of samples with various sizes (1000, 5000, 10000, 15000, and 20000) was generated from the above mentioned distributions.
2. The Fast cICA model was applied to the dataset to obtain the de-mixing matrix *W*.
3. The circular cICA model was applied to the data to learn the de-mixing matrix *W*, starting from a random complex matrix.
4. The phase-aware cICA model was applied to the data to learn the de-mixing matrix *W*.

If the source signals are perfectly separated by these algorithms, the product of the mixing matrix, *A*, and the estimated transformation *W* must be a permutation of the identity matrix. This matrix *P* = *WA* is called a performance matrix. Examples of the performance matrices *P* obtained from the fast cICA, circular and phase-aware cICA models are shown in Fig. 2.A. The performance matrix of the phase-aware cICA model is closer to the permutation of identity matrix than the performance matrix of the circular cICA model, which indicates that the phase-aware cICA performs better in source separation. To evaluate the quality of the separation, the Amari index [24] was calculated:

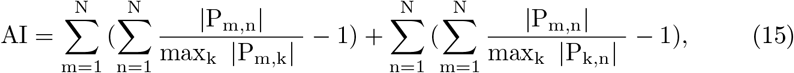

where *P*_*m,n*_ is a (m,n)-element of the performance matrix *P*. It quantifies how close the performance matrix is to a permutation of the identity matrix. The lower Amari index indicates better quality in separation (zero for perfect separation).

**Figure 2:**
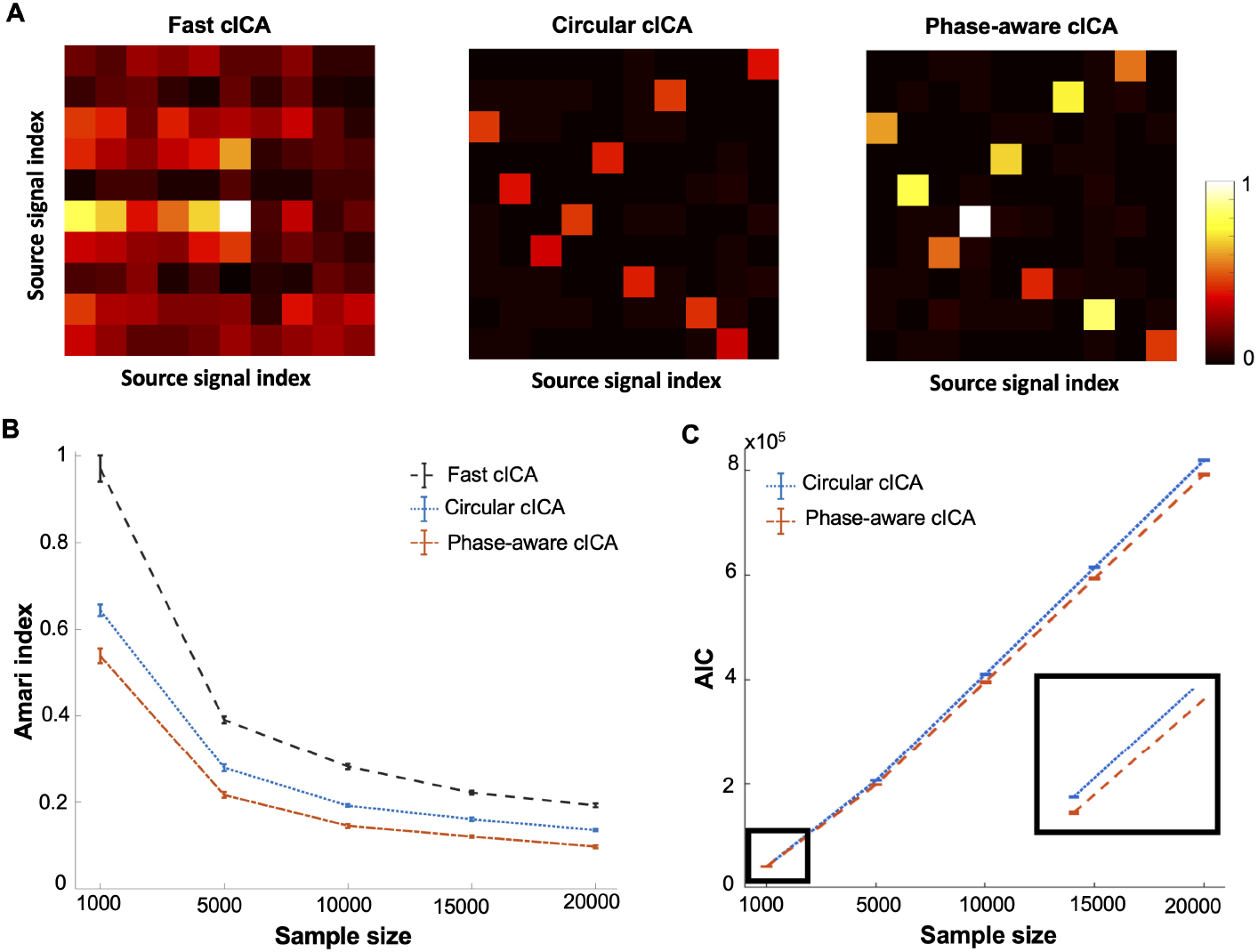
Model’s performance of Fast cICA, circular cICA and phase-aware cICA models. A) The performance matrix was computed as a product of a mixing matrix and a de-mixing matrix estimated by the three proposed algorithms (Left: Fast cICA, Middle: circular cICA Right: phase-aware cICA). The data were generated from a mixtures of 10 independent complex-valued signals, using a non-Uniform phase distribution. The performance matrices are normalized by the absolute maximum value in each column. The performance matrix of the phase-aware model is closer to the permutation of the identity matrix. This comparison shows that the phase-aware cICA performs better than the Fast cICA and circular cICA. B) The Amari indexes of the models are plotted as a function of the sample size (a mean of 20 data sets(± SD). The complex source signals were sampled from a non-uniform phase distribution (see Figure A1 for results of uniformly distributed phase signals). Zero values of the Amari index indicates the best performance of the model in separation of source signals. C) The Akaike information criterion (AIC) of both circular cICA and phase-aware cICA models were calculated from the likelihood values. This comparison exhibits a better representation of signals by the phase-aware model for all sample sizes.

The separation was tested using the circular cICA, phase-aware cICA, and Fast cICA [22]. Figure 2.B shows performance of the models for separating mixed independent complex-valued signals when they are applied to mixed complex signals generated by a combination of the bimodal and uniform phase distributions. Overall, the performance of all three models increases with the sample sizes, and the performance of the phase-aware cICA (the red plot) is the best among the three (the lowest Amari index). Model selection by Akaike information criteria also supports the better performance of the phase-aware cICA (Fig. 2.C). When applied to mixed complex signals generated by the uniform phase distribution (i.e., circular complex variables), the circular and phase-aware cICAs outperformed the Fast complex ICA (Supplementary Figure 1). Importantly, the performance of the phase-aware cICA approaches that of circular cICA whose assumption coincides with the data, indicating that the phase-aware cICA can successfully estimate the uniform phase distribution in the data.

### Application of phase-aware cICA to natural scenes

In this section, we apply the phase-aware cICA model to natural scenes. We used the Hans van Hateren’s repository of natural scenes [25] provided by Olshausen and Field [26]. We randomly selected 50, 000 image patches with size 10 × 10 pixels from the natural scenes. We then computed the Fourier transform of each patch, and obtained the complex representation of natural scenes. After the mean values of each complex-valued patches were subtracted, we performed the complex whitening algorithm on the data, based on the complex principle component analysis (complex PCA). See Appendix 5 for details of the FFT and complex PCA. Finally, we applied the algorithms (circular cICA and phase-aware cICA) to the whitened natural patches to obtain the complex-valued features and source signals. In Appendix 5, we also explain how this procedure constructs a generative model of the real-valued image patches that imposes structured amplitudes and phases in its complex representation. For comparison, we also analyzed the natural scences by the Fast cICA.

To achieve convergence on learning as quickly as possible, we used the Adam optimiser for the gradient search of Eq. 7 and 12 as implemented in [27] with the suggested default parameters and a suitable initial learning rate. Further, learning of the phase-aware model could be accelerated if we start from the converged features of the circular cICA, and then simultaneously learn the features and shape parameters, *κ*_*i*_ (*i* = 1, …, *N*). However, it turns out that the gradient of the shape parameters is zero if we start learning the phase-aware model from the uniform phase (*κ*_*i*_ = 0). The stability analysis (Appendix 4) reveals that the uniform phase distribution is a local minimum if sample phases satisfy

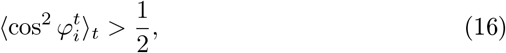

where *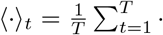·*. Because cos^2^ *φ*_*i*_ has a peak at *φ*_*i*_ = 0 or *π*, this means that the model of a uniform phase distribution (*κ*_*i*_ = 0) is a local minimum (the expectation is larger than 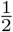) if the empirical distribution of 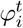 is concentrated within 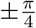 of the phase 0 or *π*. To the contrary, if 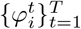 are concentrated outside of these ranges, the model with a uniform phase distribution is a local maximum.

Following this stability analysis, we propose to add independent weak positive noise sampled from a distribution such as a uniform or Gamma distribution to *κ*_*i*_ (*i* = 1, …, *N*). This makes the gradients non-vanishing, and leads to the optimal *κ* according to the distribution of 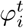of the source signals obtained during learning. Accordingly, we optimize the phase-aware model as follows:

1. We first run the circular cICA optimization until it converges to obtain the optimal values of *β* and **W**.
2. We optimize the phase-aware cICA with initial values given by the circular cICA, while initial values of *κ* are sampled from an uniform distribution [0, *ϵ*], where *ϵ* is a small value.

The likelihood increase is shown in Figure 3. This improvement is significant (p < 10^−3^) by the likelihood-ratio test with the degree of freedom *N* (i.e., dimension of the shape parameters, 100 in our case). The significant improvement of the goodness-of-fit by the phase-aware model indicates that structured spatial phase information is important in the redundancy reduction of natural scences.

**Figure 3:**
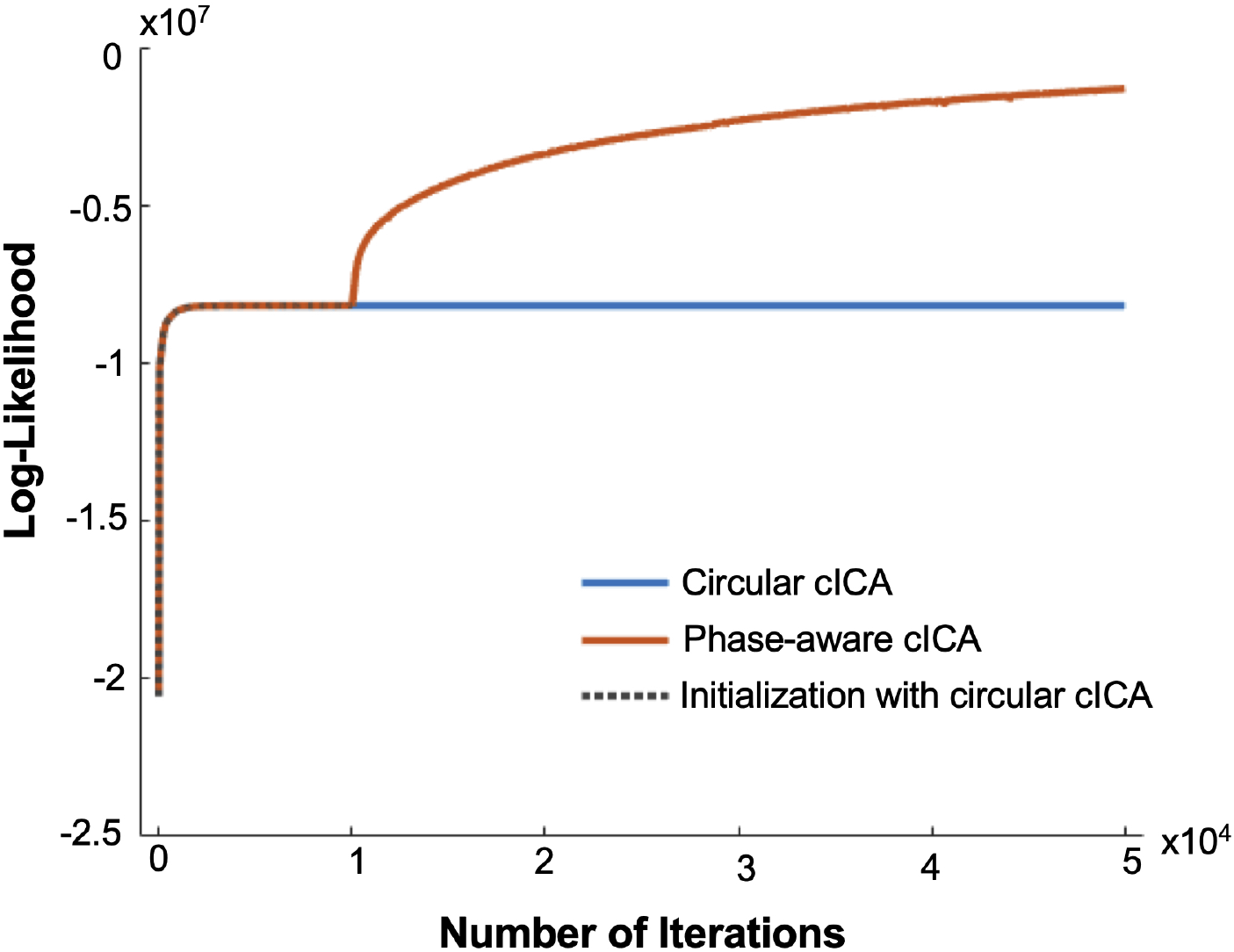
Comparison of log-likelihood functions for the circular cICA and phase-aware cICA when initialized with the circular cICA optima

Figure 4 shows the features extracted from the natural scenes. In order to show the features for real-valued images, we show the complex features obtained after performing inverse of the complex ICA and then the inverse Fourier transform to the learned de-mixing matrix *W* (See Appendix 5). Both the real and imaginary parts are shown. As a comparison, we show the features obtained from the Fast cICA applied to the same natural patches. By comparing the complex features obtained from the Fast cICA and phase-aware cICA (Figure 4), we conclude that the phase-aware cICA model provides features that are more close to the receptive fields of neurons in early visual cortex than the Fast cICA model. Further, many pairs of the components are in quadrature phases, which could explain observed topographic relations between nearby simple cells [19].

**Figure 4:**
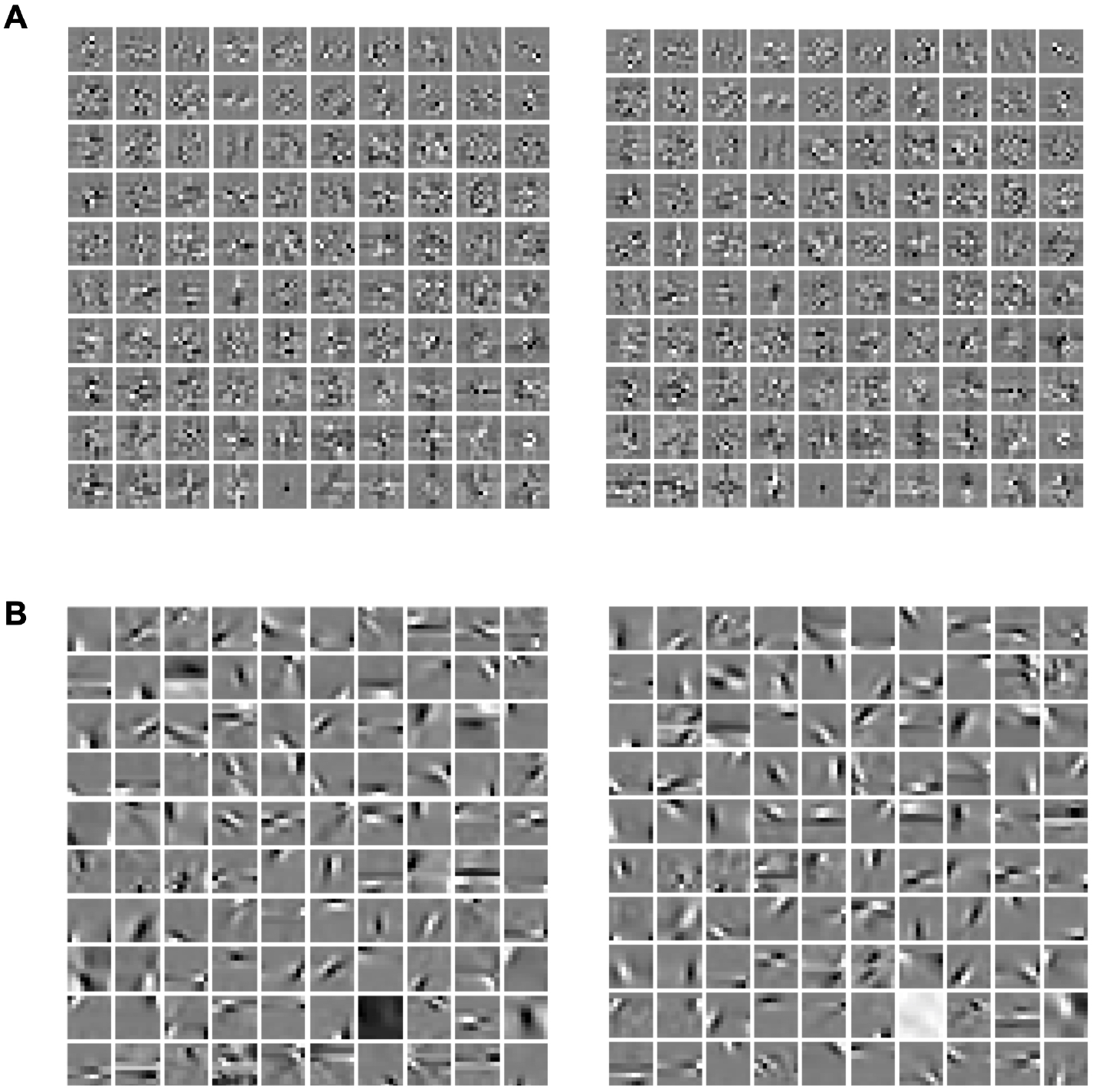
Complex features learned from natural scenes. The whole set of Real (Left) and Imaginary (Right) components of complex features obtained by the Fast cICA and phase-aware cICA are represented in the panel A and B, respectively. Both models were applied to the patches with 10*10 pixels randomly selected from natural images. The complex basis functions of phase-aware cICA are very close with the receptive field of neurons observed in V1 area of early visual cortex.

## 4. Conclusion

In this study, we introduced a generative model for complex representation of natural scenes, which is generally applicable for separation of non-circular complex source signals. We demonstrated that, in blind source separation of complex-valued signals, the proposed model outperforms the other methods that do not consider the phase information because the proposed model adaptively infers the underlying phase distribution. Applied to the natural scenes, we found that the components of learned complex feature are similar to Gabor filters, and that some pairs of components had quadrature phases. These results are consistent with proposals in signal processing to use quadrature pairs of Gabor filters [28, 29]. Under the efficient coding hypothesis, it explains characteristic of the simple-cell receptive fields and observed topographic relations between nearby simple cells [19]. In sum, superior performance of the phase-aware model and the efficient coding hypothesis suggest that visual systems are adapted to model spatial phase statistics in natural scenes.

## 5. Acknowledgment

This work was supported in part by the Cooperative Intelligence Joint Research between Kyoto University and Honda Research Institute Japan. HM acknowledges financial supports of IPM and RIKEN BSI during his PhD study. HM was supported by the EPSRC Program (Grant No. EP/P006094/1: Brains on Board).

