## APPENDIX for "Learning Complex Representations from Spatial Phase Statistics of Natural Scenes"

The appendices 1-4 contain derivations of gradients for the maximum likelihood of the proposed models. In Appendix 1, we provide definitions of derivatives for the complex values and functions. The gradient of circular complex-valued ICA (cICA) is computed in Appendix 2. The gradient of phase-aware cICA is given in Appendix 3. In Appendix 5, we introduce procedure to analyze real-valued signals by the proposed methods.

#### Appendix 1: Wirtinger calculus

Let $z$ be a complex value given by $z = x+jy$, and $z^*$ be its conjugate ($z^* = x-jy$). For the expressions involving derivatives of complex quantities, we use 'Wirtinger derivatives' (1; 2):

$$\frac{\partial}{\partial z} := \frac{1}{2}[\frac{\partial}{\partial x} - j\frac{\partial}{\partial y}], \tag{1}$$

and

$$\frac{\partial}{\partial z^*} := \frac{1}{2}[\frac{\partial}{\partial x} + j\frac{\partial}{\partial y}], \tag{2}$$

Using these expressions, the following identity is obtained:

$$\frac{\partial z^*}{\partial z} = \frac{\partial z}{\partial z^*} = 0, \tag{3}$$

From Eq. 3, we see that $z$ and $z^*$ can be treated as constants with respect to each other while performing differentiation. Further we have

$$\frac{\partial |z|^2}{\partial z} = \frac{\partial}{\partial z} zz^* = z^*, \tag{4}$$

and

$$\frac{\partial |z|}{\partial z} = \frac{1}{2|z|}\frac{\partial |z|^2}{\partial z} = \frac{z^*}{2|z|}. \tag{5}$$

These relations will be used in the subsequent calculations.

The differential $df$ for a differentiable complex-valued function $f$ on complex domains is written using $z$ and $z^*$ as independent variables:

$$df = \frac{\partial f}{\partial z}dz + \frac{\partial f}{\partial z^*}dz^*, \tag{6}$$

where the complex derivatives are given by Eq. 1 and Eq. 2. For a real-valued function including the likelihood function, we have $\frac{\partial f}{\partial z}dz = \left(\frac{\partial f}{\partial z^*}dz^*\right)^*$ because $f^* = f$. Hence, Eq. 6 simplifies to

$$df = 2\,\mathrm{Re}\left(\frac{\partial \mathrm{f}}{\partial \mathrm{z}}\mathrm{dz}\right). \tag{7}$$

To find the direction of steepest ascent, we have to maximize $df$. We note that the differential is bounded as

$$df = 2\,\mathrm{Re}\left(\frac{\partial \mathrm{f}}{\partial \mathrm{z}}\mathrm{dz}\right) \leq 2\left|\frac{\partial \mathrm{f}}{\partial \mathrm{z}}\mathrm{dz}\right|,$$

with equality holding only if $\frac{\partial f}{\partial z}dz$ is real. This condition is realized when $dz$ is a conjugate of $\frac{\partial f}{\partial z}$: $dz = \left(\frac{\partial f}{\partial z}\right)^* = \frac{\partial f}{\partial z^*}$. Hence, the update equation of $z$ for a real cost function $f$ on the complex domain is written as

$$z \leftarrow z + 2k\frac{\partial f}{\partial z^*},$$

where $k$ is the learning coefficient. See section 2.3.10 of (3) for further details on the derivatives of complex values and functions.

### Appendix 2: Gradient of circular complex-valued ICA

In this appendix, we derive gradients of the likelihood function for the circular complex-valued ICA with respect to its parameters. The likelihood function was obtained as

$$l(\boldsymbol{W},\ \boldsymbol{\beta};\ \boldsymbol{X}^{\mathbf{obs}}) = \sum_{t=1}^{T}\sum_{i=1}^{N}[2\log\ \beta_i\ - \beta_i r_i^t] - TN\log\ 2\ \pi + T\log\det\overline{\boldsymbol{W}}, \tag{8}$$

where $\boldsymbol{W} \in C^{N\times N}$ and $\boldsymbol{\beta} \in R^{N\times 1}$. We will derive the gradients of the likelihood function defined above in Eq. 8 with respect to $\boldsymbol{W}$ and $\boldsymbol{\beta}$.

First, we obtain its derivatives with respect to $\beta_i$ $(i = 1, \dots, N)$:

$$\frac{\partial l(\boldsymbol{W},\ \boldsymbol{\beta})}{\partial \beta_i} = \frac{\partial}{\partial \beta_i}\sum_{t=1}^{T}\sum_{i=1}^{N}[2\log\ \beta_i\ - \beta_i r_i^t]. \tag{9}$$

This equation was obtained because the term $\overline{\boldsymbol{W}}$ does not depend on $\boldsymbol{\beta}$, and $TN\log\ 2\pi$ is a constant. The above expression can be further simplified to

$$\frac{\partial l(\boldsymbol{W},\ \boldsymbol{\beta})}{\partial \beta_i} = \sum_{t=1}^{T}(\frac{2}{\beta_i} - r_i^t). \tag{10}$$

Next, we obtain the derivatives with respect to $W_{m,n}$,

$$\frac{\partial l(W,\ \beta)}{\partial W_{m,n}} = \frac{\partial}{\partial W_{m,n}}\left(\sum_{t=1}^{T}\sum_{i=1}^{N}[-\beta_i r_i^t] + T\log\det\overline{\boldsymbol{W}}\right). \tag{11}$$

In this equation, the terms which are not related to $W_{m,n}$ were dropped. We now evaluate derivatives of the first and second term separately. The first term becomes

$$\frac{\partial}{\partial W_{m,n}}\sum_{t=1}^{T}\sum_{i=1}^{N}[-\beta_i r_i^t] = -\sum_{t=1}^{T}\sum_{i=1}^{N}\beta_i\frac{\partial r_i^t}{\partial W_{m,n}}. \tag{12}$$

Using the relation: $s_i^t = \sum_{k=1}^{N} W_{i,k}X_k^t$ (from $S = WX$) and a chain rule of derivatives, we obtain

$$\begin{aligned} -\sum_{t=1}^{T}\sum_{i=1}^{N}\beta_i\frac{\partial r_i^t}{\partial W_{m,n}} &= -\sum_{t=1}^{T}\sum_{i=1}^{N}\beta_i\frac{\partial r_i^t}{\partial s_i^t}\frac{\partial s_i^t}{\partial W_{m,n}} \\ &= -\sum_{t=1}^{T}\sum_{i=1}^{N}\beta_i\frac{\partial r_i^t}{\partial s_i^t}\frac{\partial}{\partial W_{m,n}}\sum_{k=1}^{N}W_{i,k}X_k^t. \end{aligned} \tag{13}$$

Note that $r_i^t = \left|s_i^t\right|$. Hence, using Eq. 5, and noting that the only term remaining in the summation corresponds to $i = m$ and $k = n$, one can rewrite the above as

$$-\sum_{t=1}^{T}\sum_{i=1}^{N}\beta_i\frac{\partial r_i^t}{\partial s_i^t}\frac{\partial}{\partial W_{m,n}}\sum_{k=1}^{N}W_{i,k}X_k^t = -\sum_{t=1}^{T}\frac{\beta_m {s_m^t}^* X_n^t}{2r_m^t}. \tag{14}$$

Next, we evaluate the second term in Eq. 11:

$$\frac{\partial}{\partial W_{m,n}} T \log \det \overline{\boldsymbol{W}} = T \left[ \frac{\partial}{\partial \boldsymbol{W}} \log \det \overline{\boldsymbol{W}} \right]_{m,n}. \tag{15}$$

Here $\overline{\boldsymbol{W}}$ is defined as

$$\overline{\boldsymbol{W}} = \begin{bmatrix} \mathrm{Re}(\mathbf{W}) & -\mathrm{Im}(\mathbf{W}) \\ \mathrm{Im}(\mathbf{W}) & \mathrm{Re}(\mathbf{W}) \end{bmatrix} = \begin{bmatrix} \frac{\mathbf{W}+\mathbf{W}^*}{2} & \frac{\mathbf{W}^*-\mathbf{W}}{2j} \\ \frac{\mathbf{W}-\mathbf{W}^*}{2j} & \frac{\mathbf{W}+\mathbf{W}^*}{2} \end{bmatrix}, \tag{16}$$

and it can be factorized as $\overline{\boldsymbol{W}} = \boldsymbol{A}\boldsymbol{W_p}\boldsymbol{A^{-1}}$, where

$$\boldsymbol{A} = \frac{1}{2} \begin{bmatrix} j\mathbf{I} & -j\mathbf{I} \\ \mathbf{I} & \mathbf{I} \end{bmatrix}, \tag{17}$$

Here, $\mathbf{I}$ is the identity matrix of size $N \times N$, and

$$\boldsymbol{W_p} = \begin{bmatrix} \boldsymbol{W} & 0 \\ 0 & \boldsymbol{W}^* \end{bmatrix}. \tag{18}$$

Thus, $\det \overline{\boldsymbol{W}} = \det \mathbf{W} \det \mathbf{W}^*$ and $\log \det \overline{\boldsymbol{W}} = \log \det \mathbf{W} + \log \det \mathbf{W}^*$. Therefore, we obtain

$$\left[ \frac{\partial}{\partial \boldsymbol{W}} T \log \det \overline{\boldsymbol{W}} \right]_{m,n} = T \left[ \frac{\partial}{\partial \boldsymbol{W}} \big( \log \det \mathbf{W} + \log \det \mathbf{W}^* \big) \right]_{m,n}. \tag{19}$$

We note the following matrix identities from chapter two in (4),

$$\big(\det \mathbf{W}\big)^* = \det \mathbf{W}^*, \tag{20}$$

$$\frac{\partial \det \mathbf{W}}{\partial \boldsymbol{W}} = \det \mathbf{W} \left(\mathbf{W}^{-1}\right)^{\mathrm{T}}, \tag{21}$$

From Eq. 3, $\boldsymbol{W^*}$ does not depend on $\boldsymbol{W}$. Furthermore, using Eq. 21, we can simplify Eq. 19 as

$$\left[ \frac{\partial}{\partial \boldsymbol{W}} \big( \log \det \mathbf{W} \big) \right]_{m,n} = \frac{1}{\det \mathbf{W}} \frac{\partial \det \mathbf{W}}{\partial \boldsymbol{W}} = \left(\boldsymbol{W}^{-1}\right)^{T}. \tag{22}$$

Using the expressions derived in Eq. 14 and Eq. 22, we can rewrite Eq. 11 as

$$\frac{\partial l(\boldsymbol{W},\ \boldsymbol{\beta})}{\partial W_{m,n}} = T \left(\boldsymbol{W}^{-1}\right)^{T}_{m,n} - \sum_{t=1}^{T} \frac{\beta_m X_n^t {s_m^t}^*}{2 r_m^t}, \tag{23}$$

where the superscript $T$ indicates a transpose of the de-mixing matrix, and the superscripts $*$ denotes the conjugate of complex coefficient, $s_m$. Hence, for the update, the gradient with respect to $W^*_{m,n}$ is

$$\frac{\partial l(\boldsymbol{W},\ \boldsymbol{\beta})}{\partial W^*_{m,n}} = T \left(\boldsymbol{W}^{-1}\right)^{H}_{m,n} - \sum_{t=1}^{T} \frac{\beta_m {X_n^t}^* s_m^t}{2 r_m^t}, \tag{24}$$

where $H$ denotes the hermitian (conjugate transpose) of de-mixing matrix $\boldsymbol{W}$, respectively. Note that $r_i$ and $s_m$, are calculated from $\boldsymbol{W}$ and $\boldsymbol{X}^{\mathbf{obs}}$.

### Appendix 3: Gradient of the phase-aware complex-valued ICA

In this Appendix, we derive gradients for the phase-aware cICA model. In this model, the phase distribution is modeled as a mixture of uniform and two von-Mises distributions:

$$p_{\varphi_i}(\varphi_i;\kappa_i,\lambda) = \lambda\ \mathrm{vM}(\varphi_\mathrm{i};\kappa_\mathrm{i},0) + \lambda\ \mathrm{vM}(\varphi_\mathrm{i};\kappa_\mathrm{i},\pi) + (1-2\lambda)\frac{1}{2\pi}. \tag{25}$$

Here $\mathrm{vM}(\varphi_\mathrm{i};\kappa_\mathrm{i},\mu_\mathrm{i})$ is a von-Mises distribution for a circular variable $\varphi_i$ with mean $\mu_i$ and a concentration parameter $\kappa_i$,

$$\mathrm{vM}(\varphi_\mathrm{i};\kappa_\mathrm{i},\mu_\mathrm{i}) = \frac{\mathrm{e}^{\kappa_\mathrm{i}\cos(\varphi_\mathrm{i}-\mu_\mathrm{i})}}{2\pi \mathrm{I}_0(\kappa_\mathrm{i})},$$

where $I_0(\cdot)$ is the modified Bessel function of order 0. Thus the above model in Eq. 25 exhibits bimodal structure with peaks at 0 and $\pi$, on top of the constant baseline. For simplicity we further assume equal contributions from each component ($\lambda = 1/3$). Then the phase distribution that can cover a uniform and a spectrum of bimodal phase distributions is simplified as

$$p_{\varphi_i}(\varphi_i;\kappa_i) = \frac{1}{3\pi\ I_0(\kappa_i)}\cosh(\kappa_i\ \cos\varphi_i) + \frac{1}{6\pi}. \tag{26}$$

This model is a modification of the previous circular cICA. We call this new model the phase-aware cICA.

The log-likelihood function of the phase-aware cICA model is

$$l(\boldsymbol{\theta};\boldsymbol{X}^{\mathbf{obs}}) = \sum_{t=1}^{T}\sum_{i=1}^{N}[2\log\beta_i\ - \beta_i r_i^t + \log\left(\frac{1}{3\pi\ I_0(\kappa_i)}\cosh(\kappa_i\ \cos\varphi_i^t) + \frac{1}{6\pi}\right)] + T\log\det\overline{\boldsymbol{W}}, \tag{27}$$

where $\boldsymbol{\theta} = (\boldsymbol{W},\boldsymbol{\kappa},\boldsymbol{\beta})$ is a set of the model parameters. Using Eq. 10 for the gradient with respect to $\boldsymbol{\beta}$, we derive the gradients with respect to $\boldsymbol{\kappa}$ and $\boldsymbol{W}$ here.

The gradient of the log-likelihood function with respect to $\kappa_i$ is given as follows,

$$\begin{aligned}
\frac{\partial l(\boldsymbol{\theta})}{\partial\kappa_i} &= \frac{\partial}{\partial\kappa_i}\sum_{t=1}^{T}\sum_{i=1}^{N}\log\left(\frac{1}{3\pi\ I_0(\kappa_i)}\cosh(\kappa_i\ \cos\varphi_i^t) + \frac{1}{6\pi}\right)\\
&= \sum_{t=1}^{T}\frac{\left(\frac{\cosh(\kappa_i\ \cos\varphi_i^t)}{3\pi\ I_0(\kappa_i)} + \frac{1}{6\pi}\right)'}{\frac{\cosh(\kappa_i\ \cos\varphi_i^t)}{3\pi\ I_0(\kappa_i)} + \frac{1}{6\pi}}\\
&= \sum_{t=1}^{T}\frac{1}{3\pi}\frac{\sinh(\kappa_i\cos\varphi_i^t)\cos\varphi_i^t\cdot\frac{1}{I_0(\kappa_i)} - \cosh(\kappa_i\cos\varphi_i^t)\cdot\frac{I_1(\kappa_i)}{I_0(\kappa_i)^2}}{\frac{\cosh(\kappa_i\ \cos\varphi_i^t)}{3\pi\ I_0(\kappa_i)} + \frac{1}{6\pi}}\\
&= \sum_{t=1}^{T}\frac{I_0(\kappa_i)\sinh(\kappa_i\cos\varphi_i^t)\cos\varphi_i^t - I_1(\kappa_i)\cosh(\kappa_i\cos\varphi_i^t)}{I_0(\kappa_i)\cosh(\kappa_i\ \cos\varphi_i^t) + \frac{I_0(\kappa_i)^2}{2}}.
\end{aligned} \tag{28}$$

Next, we compute the gradient with respect to $W_{m,n}$:

$$\frac{\partial l(\boldsymbol{\theta})}{\partial W_{m,n}} = \frac{\partial}{\partial W_{m,n}}\left(\sum_{t=1}^{T}\sum_{i=1}^{N}[-\beta_i r_i^t + \log\left(\frac{1}{3\pi\ I_0(\kappa_i)}\cosh(\kappa_i\ \cos\varphi_i^t) + \frac{1}{6\pi}\right)]+ T\log\det\overline{\boldsymbol{W}}\right). \tag{29}$$

The first and third terms are the same as those obtained for the previous circular cICA. To compute the derivative of the second term, we need to obtain:

$$\frac{\partial\varphi_i^t}{\partial W_{m,n}} = \frac{\partial\varphi_i^t}{\partial s_i^t}\frac{\partial s_i^t}{\partial W_{m,n}}. \tag{30}$$

Writing $s_i^t$ and $\varphi_i^t$ as functions of (x,y), namely $s_i^t = x + jy$ and $\varphi_i^t = \arctan(\frac{y}{x})$, we can obtain $\frac{\partial\varphi_i^t}{\partial s_i^t}$ using Eq. 1 as

$$\frac{\partial\varphi_i^t}{\partial s_i^t} = \frac{1}{2}\left[\frac{\partial}{\partial x}\arctan(\frac{y}{x}) - j\frac{\partial}{\partial y}\arctan(\frac{y}{x})\right] = \frac{1}{2}\left[\frac{-y-jx}{x^2+y^2}\right] = \frac{-j{s_i^t}^*}{2{r_i^t}^2}. \tag{31}$$

Using the expression for $\frac{\partial s_i^t}{\partial W_{m,n}}$ derived previously in Eqs. 13 and 14, we have

$$\frac{\partial\varphi_i^t}{\partial W_{m,n}} = \frac{-jX_n^t{s_m^t}^*}{2{r_m^t}^2} \tag{32}$$

Therefore, differentiating the second term in Eq. 29 gives us

$$\begin{aligned}
&\frac{\partial}{\partial W_{m,n}}\sum_{t=1}^{T}\sum_{i=1}^{N}\log\left(\frac{1}{3\pi\ I_0(\kappa_i)}\cosh(\kappa_i\ \cos\varphi_i^t) + \frac{1}{6\pi}\right)\\
&= \sum_{t=1}^{T}\sum_{i=1}^{N}\frac{\frac{1}{3\pi I_0(\kappa_i)}(\cosh(\kappa_i\cos\varphi_i^t))'}{\frac{1}{3\pi\ I_0(\kappa_i)}\cosh(\kappa_i\ \cos\varphi_i^t) + \frac{1}{6\pi}}\\
&= \sum_{t=1}^{T}\sum_{i=1}^{N}\frac{\frac{1}{3\pi I_0(\kappa_i)}\sinh(\kappa_i\cos\varphi_i^t)\kappa_i(\cos\varphi_i^t)'}{\frac{1}{3\pi\ I_0(\kappa_i)}\cosh(\kappa_i\ \cos\varphi_i^t) + \frac{1}{6\pi}}\\
&= \sum_{t=1}^{T}\sum_{i=1}^{N}\frac{\frac{-1}{3\pi I_0(\kappa_i)}\sinh(\kappa_i\cos\varphi_i^t)\kappa_i\sin\varphi_i^t\frac{\partial\varphi_i^t}{\partial W_{m,n}}}{\frac{1}{3\pi\ I_0(\kappa_i)}\cosh(\kappa_i\ \cos\varphi_i^t) + \frac{1}{6\pi}}\\
&= \sum_{t=1}^{T}\frac{\frac{1}{3\pi I_0(\kappa_m)}\sinh(\kappa_m\cos\varphi_m^t)\kappa_m\sin\varphi_m^t\frac{jX_n^t{s_m^t}^*}{2{r_m^t}^2}}{\frac{1}{3\pi\ I_0(\kappa_m)}\cosh(\kappa_m\ \cos\varphi_m^t) + \frac{1}{6\pi}},
\end{aligned}$$

where the summation with respect to $i$ turns out to be the single term for which $i = m$.

Putting it all together, we obtain the gradient as

$$\frac{\partial l(\boldsymbol{\theta})}{\partial W_{m,n}} = T(\boldsymbol{W}^{-1})^T{}_{m,n} - \sum_{t=1}^{T}[\frac{\beta_m X_n^t{s_m^t}^*}{2\ r_m^t} - \frac{\frac{1}{3\pi I_0(\kappa_m)}\sinh(\kappa_m\cos\varphi_m^t)\kappa_m\sin\varphi_m^t\frac{jX_n^t{s_m^t}^*}{2{r_m^t}^2}}{\frac{1}{3\pi\ I_0(\kappa_m)}\cosh(\kappa_m\ \cos\varphi_m^t) + \frac{1}{6\pi}}]. \tag{33}$$

Hence, for the update, the gradient with respect to $W^*_{m,n}$ becomes

$$\frac{\partial l(\boldsymbol{\theta})}{\partial W^*_{m,n}} = T(\boldsymbol{W}^{-1})^H{}_{m,n} - \sum_{t=1}^{T}[\frac{\beta_m X_n^{t\,*} s_m^t}{2\ r_m^t} + \frac{\frac{1}{3\pi I_0(\kappa_m)}\sinh(\kappa_m \cos\varphi_m^t)\kappa_m \sin\varphi_m^t \frac{j X_n^{t\,*} s_m^t}{2 {r_m^t}^2}}{\left(\frac{1}{3\pi\ I_0(\kappa_m)}\cosh(\kappa_m\ \cos\varphi_m^t) + \frac{1}{6\pi}\right)}]. \tag{34}$$

### Appendix 4: Stability analysis of the phase-aware cICA model

In this appendix, we examine stability of the phase-aware model around the circular cICA model by examining the second derivative of the likelihood function with respect to the shape parameter. Based on this analysis, we propose to perform learning of the phase-aware model from the features learned from circular model, but with weakly perturbed shape parameters.

First, let us define the numerator and denominator in Eq. 28 as

$$
\begin{aligned}
u(\kappa_i, \varphi_i^t) &= I_0(\kappa_i)\sinh(\kappa_i \cos\varphi_i^t)\cos\varphi_i^t - I_1(\kappa_i)\cosh(\kappa_i \cos\varphi_i^t), \\
v(\kappa_i, \varphi_i^t) &= I_0(\kappa_i)\cosh(\kappa_i \cos\varphi_i^t) + \frac{I_0(\kappa_i)^2}{2}.
\end{aligned}
$$

Note that we have $I_0(0) = 1$ and $I_1(0) = 0$ when $\kappa_i = 0$, from which we obtain $u(0, \varphi_i^t) = 0$ and $v(0, \varphi_i^t) = 1 + \frac{1}{2} = \frac{3}{2}$. Thus the gradient at $\kappa_i = 0$ is zero:

$$
\left.\frac{\partial l(\boldsymbol{\theta})}{\partial \kappa_i}\right|_{\kappa_i = 0} = 0. \tag{35}
$$

This means that the log likelihood function takes either a local maximum or minimum at $\kappa_i = 0$, along this dimension (Note that this does not indicate that it is also a maximum or minimum in directions for other parameters).

In order to determine whether the likelihood function exhibits a local maximum or minimum at $\kappa_i = 0$, we perform the following stability analysis. The second derivative of the log likelihood function is give as

$$
\frac{\partial l(\boldsymbol{\theta})^2}{\partial^2 \kappa_i} = \sum_{t=1}^{T} \frac{\partial}{\partial \kappa_i} \frac{u(\kappa_i, \varphi_i^t)}{v(\kappa_i, \varphi_i^t)} = \sum_{t=1}^{T} \frac{u'(\kappa_i, \varphi_i^t)v(\kappa_i, \varphi_i^t) - u(\kappa_i, \varphi_i^t)v'(\kappa_i, \varphi_i^t)}{v(\kappa_i, \varphi_i^t)^2}. \tag{36}
$$

To evaluate this equation, we compute the derivative of $u$ and $v$. First, the derivative of $u$ is given as

$$
\begin{aligned}
u'(\kappa_i, \varphi_i^t) = {} & I_0(\kappa_i)\cosh(\kappa_i \cos\varphi_i^t)\cos^2\varphi_i^t + \cancel{I_1(\kappa_i)\sinh(\kappa_i \cos\varphi_i^t)\cos\varphi_i^t} \\
& - \cancel{I_1(\kappa_i)\sinh(\kappa_i \cos\varphi_i^t)\cos\varphi_i^t} - \frac{1}{2}(I_0(\kappa_i) + I_2(\kappa_i^t))\cosh(\kappa_i \cos\varphi_i^t) \\
= {} & I_0(\kappa_i)\cosh(\kappa_i \cos\varphi_i^t)\cos^2\varphi_i^t - \frac{1}{2}(I_0(\kappa_i) + I_2(\kappa_i))\cosh(\kappa_i \cos\varphi_i^t),
\end{aligned}
$$

where we used the identity $\frac{dI_0(x)}{dx} = I_1(x)$ and $\frac{dI_1(x)}{dx} = \frac{1}{2}(I_0(x) + I_2(x))$. Next, the derivative of $v$ is

$$
v'(\kappa_i, \varphi_i^t) = I_0(\kappa_i)\sinh(\kappa_i \cos\varphi_i^t)\cos\varphi_i^t + I_1(\kappa_i)\cosh(\kappa_i \cos\varphi_i^t) + I_0(\kappa_i)I_1(\kappa_i).
$$

From these equations, we obtain $u'(0, \varphi_i^t) = \cos^2\varphi_i - \frac{1}{2}$ and $v'(0, \varphi_i^t) = 0$. Together with $u(0, \varphi_i^t) = 0$ and $v(0, \varphi_i^t) = \frac{3}{2}$, the second derivative evaluated at $\kappa_i = 0$ is

$$
\begin{aligned}
\left.\frac{\partial l(\boldsymbol{\theta})^2}{\partial^2 \kappa_i}\right|_{\kappa_i = 0} &= \sum_{t=1}^{T} \frac{\left(\cos^2\varphi_i^t - \frac{1}{2}\right)\cdot\frac{3}{2} - 0\cdot 0}{\left(\frac{3}{2}\right)^2} \\
&= \sum_{t=1}^{T} \frac{2}{3}\left(\cos^2\varphi_i^t - \frac{1}{2}\right).
\end{aligned} \tag{37}
$$

This result suggests that the circular model ($\kappa_i = 0$, an assumption of a uniform phase distribution) is a local minimum if

$$\langle \cos^2 \varphi_i^t \rangle_t > \frac{1}{2}, \tag{38}$$

where $\langle \cdot \rangle_t = \frac{1}{T} \sum_{t=1}^{T} \cdot$. Namely, the empirical expectation of $\cos^2 \varphi_i^t$ has to be greater than $\frac{1}{2}$. Note that, if observed $\varphi_i^t$s are uniformly distributed, expectation of $\cos^2 \varphi_i^t$ is given as

$$\lim_{t \to \infty} \langle \cos^2 \varphi_i^t \rangle_t = \int_0^{2\pi} \cos^2 \varphi_i \, d\varphi_i = \frac{1}{2}. \tag{39}$$

Based on the functional form of $\cos^2 \varphi_i$ whose peaks are located at the phase 0 and $\pi$, this means that the circular model ($\kappa_i = 0$) is a local minimum (the expectation is larger than $\frac{1}{2}$) if the empirical distribution of $\varphi_i^t$ is concentrated within $\pm\frac{\pi}{4}$ at the phase 0 or $\pi$. Since our phase-aware model has peaks at 0 and $\pi$ if $\kappa_i > 0$, the phase-aware model yields larger likelihood than the model with a uniform phase distribution for such data. To the contrary, if $\{\varphi_i^t\}_{t=1}^T$ are concentrated outside of these ranges, then the model with a uniform phase distribution is a local maximum: The phase-aware model yields lower likelihood than the uniform phase model for such data.

Based on the above reasoning, we select initial values of $\kappa_i$ for learning as follows. We add independent weak positive noise sampled from either Uniform or Gamma distribution to $\kappa_i$ ($i = 1, \ldots, N$), which are initially set to zeros. This makes the gradients non-vanishing, and leads to the optimal $\kappa_i$ according to the emprical distribution of $\varphi_i^t$ of the source signals obtained during learning.

### Appendix 5: Complex representation of real-valued signals

In this appendix, we explain how the real-valued signal is modeled through complex representation.

In order to analyze real-valued signals such as natural images, we first applied the fast Fourier transform (FFT) to the original real-valued signal, $X_{\text{real}}$. Let $F$ be the FFT operator. Then complex-valued representation of the real-valued data is given as $X = FX_{\text{real}}$.

In real-valued ICA, it is common to remove the second order correlations with PCA. In this article, we used the generalization of the technique to complex matrices (complex PCA). In this method, we first obtain the complex-valued covariance matrix $C = E[XX^H]$, where $X^H$ is the Hermitian (conjugate transpose), and then perform the eigen-decomposition $C = UDU^*$ to obtain the whitening matrix $Q_{pca} = \sqrt{D^{-1}}U$. Decomposition of the whitened signal $Q_{pca}X$ is obtained as

$$Q_{pca}X = A\,S. \tag{40}$$

In the main text of this article, the mixing matrix $A$, or the de-mixing matrix $W(= A^{-1})$, was learned from the whitened data. Therefore decomposition of the original complex data $X$ is obtained as $X = Q_{pca}^{-1}AS$. Namely, $Q_{pca}^{-1}A$ gives the mixing matrix that returns $X$ from $S$. Using the learned matrix $W$, this mixing matrix is given as $Q_{pca}^{-1}W^{-1} = (WQ_{pca})^{-1}$.

Using the original real-valued signal $X_{\text{real}}$, Eq. 40 is expressed as $Q_{pca}FX_{\text{real}} = AS$, from which we obtain

$$\begin{aligned} X_{\text{real}} &= F^{-1}Q_{pca}^{-1}A\,S \\ &= F^{-1}(WQ_{pca})^{-1}\,S. \end{aligned} \tag{41}$$

Here $F^{-1}$ is an inverse Fourier transform operation. Let $B^* = F^{-1}(WQ_{pca})^{-1}$ be a conjugate of the complex matrix $B$, which is a collection of the $N \times 1$ column vectors: $B = [B_1, B_2, \ldots, B_N]$. Here the $i$th column is further represented as $B_i = B_i^{\text{Re}} + jB_i^{\text{Im}}$. Similarly, the element of $S$ is ginven as $s_i = s_i^{\text{Re}} + js_i^{\text{Im}}$. Then Eq. 41 is written as

$$\begin{aligned} X_{\text{real}} = B^*\,S &= \sum_{i=1}^{N} B_i^*\,s_i \\ &= \sum_{i=1}^{N}(B_i^{\text{Re}} - jB_i^{\text{Im}})(s_i^{\text{Re}} + js_i^{\text{Im}}). \end{aligned} \tag{42}$$

Under the assumption that our structured model regarding the complex sources $S$ successfully captures the statistical structure of real-valued signal, it is expected that the right-hand side of Eq. 42 is also the real-valued vector, namely:

$$X_{\text{real}} \approx \sum_{i=1}^{N}(s_i^{\text{Re}}B_i^{\text{Re}} + s_i^{\text{Im}}B_i^{\text{Im}}). \tag{43}$$

Eq. 43 constitutes a generative model of the real-valued signals. Since we impose structured dependency between the real and imaginary parts of the complex coefficients through modeling their amplitude and phase, this approximate model belongs to a class of bilinear models (5).

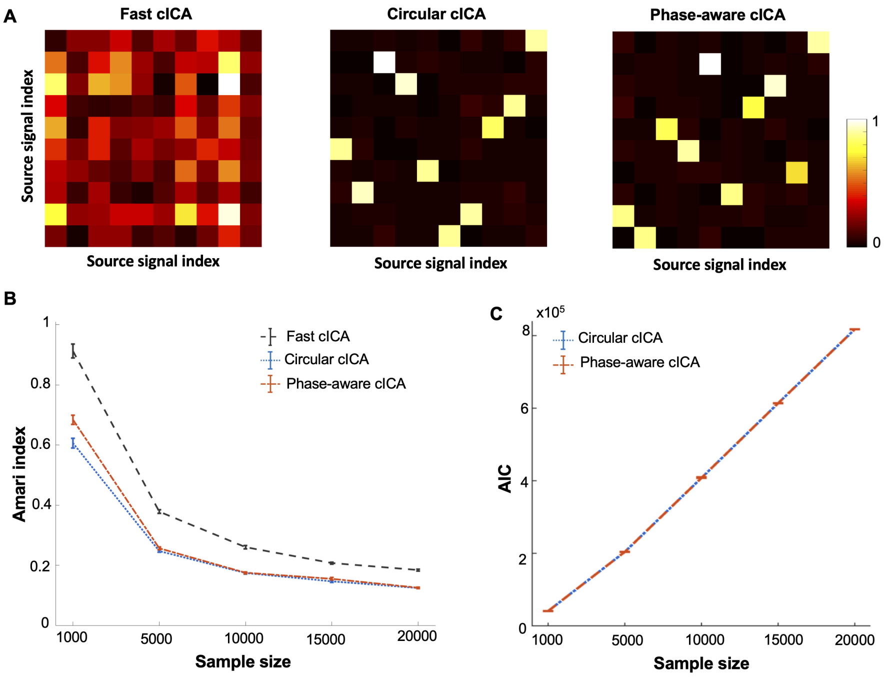


Figure 1: Performance of the models when complex source signals are sampled from a uniform phase distribution. Notations in the figure follow Fig. 2 in the main text. The same performance of the phase- aware cICA and circular cICA in separation of complex source signals with the Uniform phase assumption indicates that the phase-aware cICA can successfully estimate the uniform phase distribution in the data.
